# Automatic OpenAPI to Bio.tools Conversion

**DOI:** 10.1101/170274

**Authors:** Egon Willighagen, Jonathan Mélius

## Abstract

Computation has become a central component of life sciences research. Making computational services FAIR has had a strong interest from the life sciences community in the past 15 years. Admittedly, uptake of any of the developed solutions has been limited, and the existence of multiple approaches will not have helped. Interoperability of solution may be essential. This work introduces an interoperability layer between two approaches for FAIR annotation of web services: OpenAPI and bio. tools.

## Introduction

For many years finding webservices and documenting webservice functionality has been a key topic in Europe. For example, projects like EMBRACE and BioCatalogue annotated SOAP webservices [Rice2006, Bhagat2010]. Recently, ELIXIR started the bio.tools service [Ison2016], with a broader scope and no longer limited to SOAP webservices. With alternatives being proposed, such as the self-documenting XMPP web services [Wagener2009] and SADI [Wilkinson2011], this is welcome. Similarly, REST [Fielding2000] and REST-like webservices are starting to get documented and annotated with OpenAPI (https://www.openapis.org/), previously called Swagger. OpenAPI has the same goal as bio.tools, as well as other annotation solutions, such as BioSchemas [Larcombe2017]: FAIR annotation [Wilkinson2016, Mons2017] of webservices. There is increasing notion that this is needed to move the field forward [Exner2017, Hanwell2017]. Therefore, being able to interconvert specifications is a useful interoperability feature.

To implement this interoperability layer, ELIXIR-DK and Maastricht University set up a small project, focusing on converting OpenAPI 2.0 documentation file into bio.tools information. One point of interest here is that bio.tools support ontological annotation (using the EDAM ontology [Ison2013]), something that is not availlable in OpenAPI 2.0. This document reports on the results of this project.

## Methods

### OpenAPIs for existing REST services

To demonstrate the potential impact of this work, we manually defined an OpenAPI 2.0 JSON specification file of an existing REST interface, namely that of the Ensembl database [Flicek2013]. The online Ensembl API information was used to identify the various methods to be made available in the OpenAPI JSON.

### Crowdsourcing OpenAPIs for the life sciences

In autumn 2016, Twitter and Biostars were used to interest people in crowdsourcing and provide information about webservices in the life sciences with OpenAPIs [Willighagen2016a,Willighagen2016b]. It was decided to record the OpenAPI in Wikidata [Mietchen2015], where existing statement models were available to record “URL”s for databases, and the “instance of” qualifier was used to indicate that the URL represented a OpenAPI endpoint. The following model was selected to record the availability APIs (alongside an example annotation of a SPARQL end point):

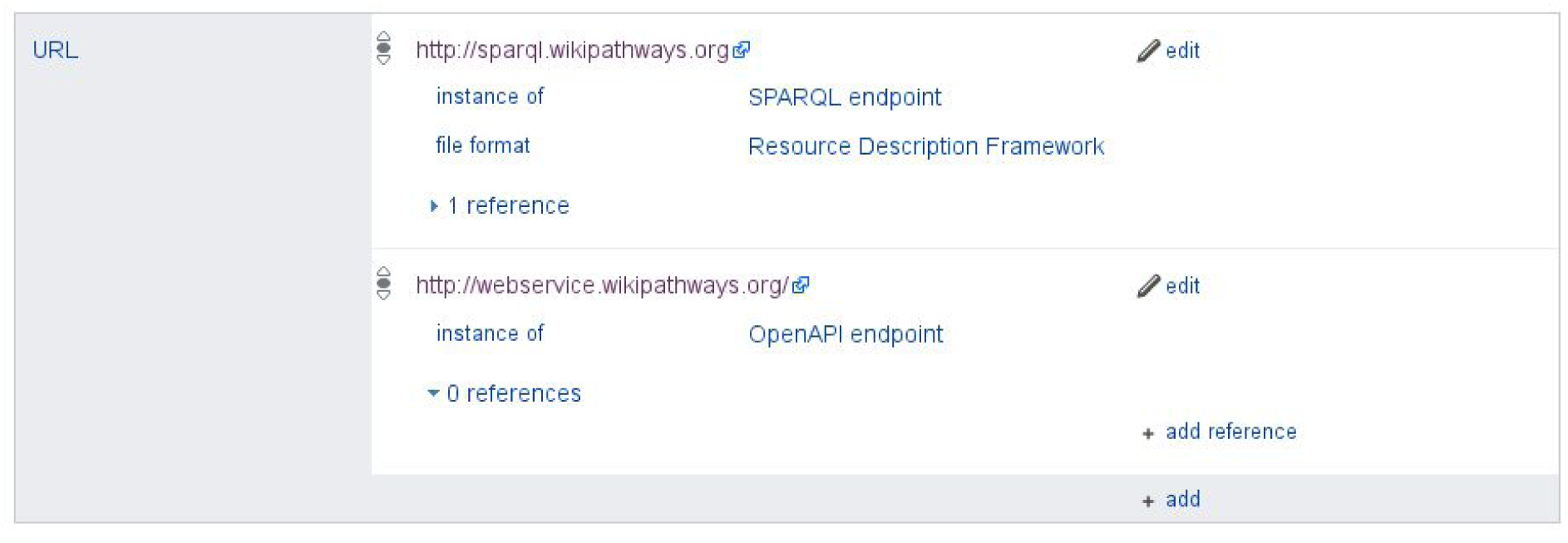

These can be recovered from Wikidata with the following SPARQL query:

~~~
SELECT ?database ?databaseLabel ?license ?licenseLabel ?value
WHERE {
 ?database ?p ?wds.
 OPTIONAL { ?database wdt:P275 ?license }
 ?wds ?v ?value.
 ?wdP wikibase:statementProperty ?v.
 ?wdP wikibase:claim ?p.
 ?wds pq:P31 wd:Q27075870.
 SERVICE wikibase:label {
  bd:serviceParam wikibase:language "en".
 }
} ORDER BY ASC(?databaseLabel)
~~~

This query can be easily repeated using <u>this</u> <u>link</u>. The resulting table shows the database entry in Wikidata, name of the database, the license of the data (entry and name), and the location of the OpenAPI endpoint.

This annotation in Wikidata was set up to avoid having to register these services in bio.tools manually, and to allow the conversion tool to be used for this.

### JSON for ontology annotation

To overcome the discrepancy in the OpenAPI JSON with respect to the ontology annotation, the concept of an external annotation file was set up, also in the JSON format. For the annotation the EDAM ontology is used [Ison2013].

### The convertor

The Java programming language was used to implement the convertor. The convertor is designed to support the OpenAPI 2.0 specification and has been tested by applying it to a number of OpenAPI 2.0 webservices. The tool can be compiled using the Maven build system.

## Results

The following sections shows the results of our efforts to show how community OpenAPI service documentation can be used in a semi-automated way to populate bio.tools. We first report on the OpenAPI JSON we created for a key European resource for the life sciences, Ensembl. We then show a, not too well succeeded, crowdsourcing of OpenAPI specifications of life sciences-related webservices. This is followed by results showing our annotation with EDAM ontology terms, and conversion to bio.tools input. We conclude with reporting on new and update bio.tools entries, enriched with specific API calls.

### Wikidata OpenAPI listing

The crowdsourcing of OpenAPIs for the live sciences yielded 14 OpenAPI-based web services related to the life sciences, as given in the below table. This table was generated using an appropriate SPARQL query.

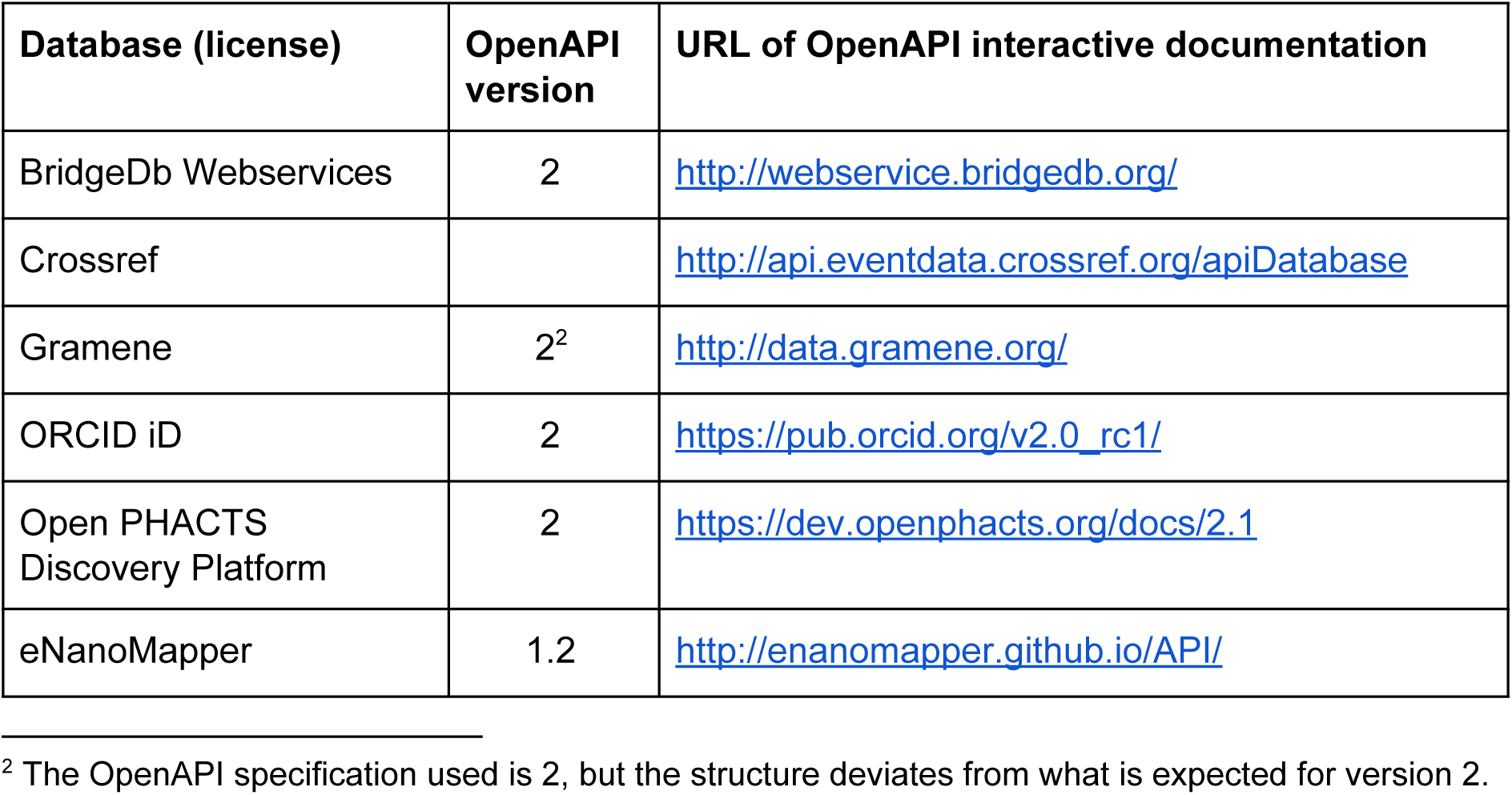
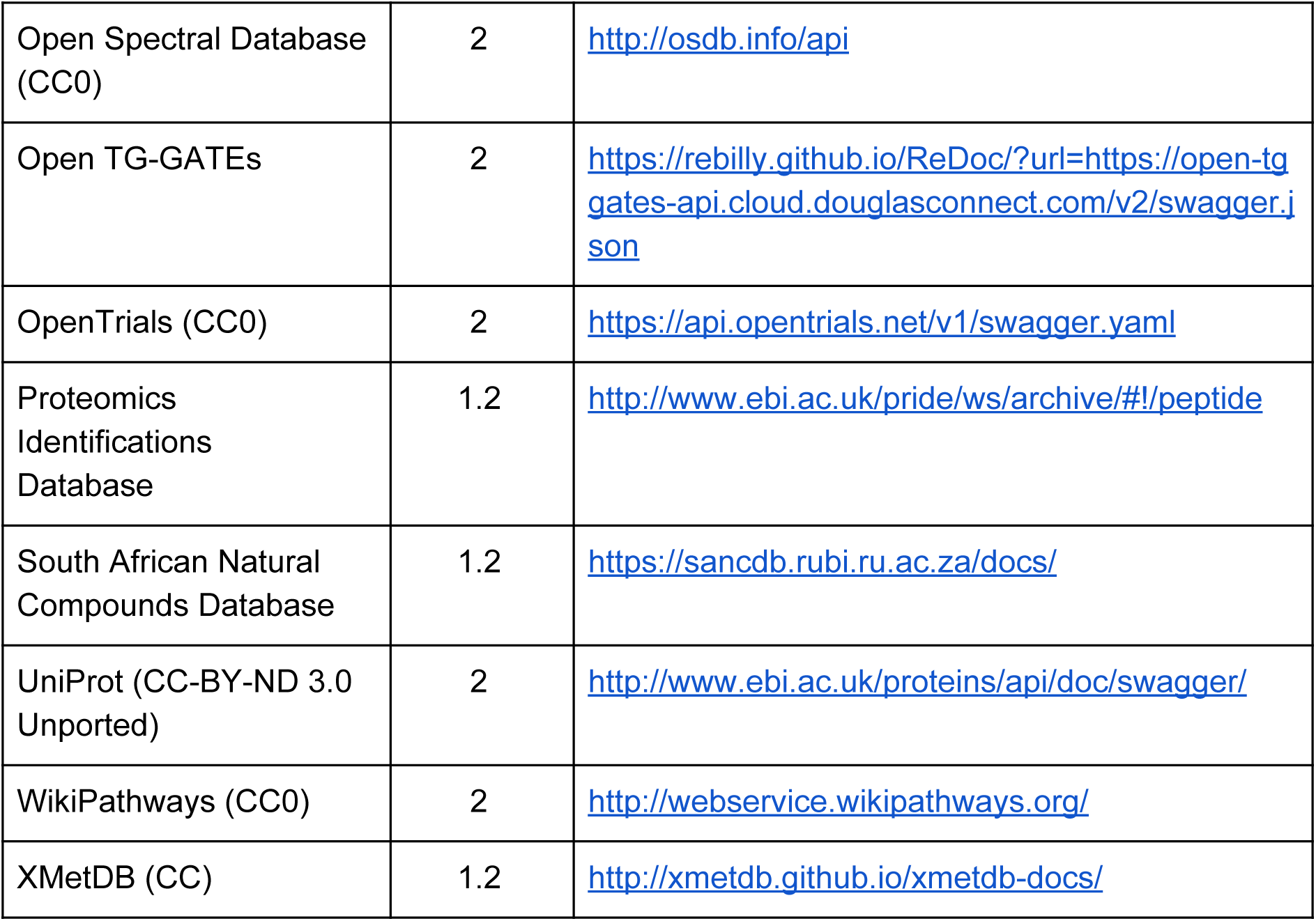

| Database (license) | OpenAPI version | URL of OpenAPI interactive documentation |
| --- | --- | --- |
| BridgeDb Webservices | 2 | http://webservice.bridgedb.org/ |
| Crossref |  | http://api.eventdata.crossref.org/apiDatabase |
| Gramene | 2<sup>2</sup> | http://data.gramene.org/ |
| ORCID iD | 2 | https://pub.orcid.org/v2.0_rc1/ |
| Open PHACTS Discovery Platform | 2 | https://dev.openphacts.org/docs/2.1 |
| eNanoMapper | 1.2 | http://enanomapper.github.io/API/ |
| Open Spectral Database (CC0) | 2 | http://osdb.info/api |
| Open TG-GATEs | 2 | https://rebilly.github.io/ReDoc/?url=https://open-tg-gates-api.cloud.douglasconnect.com/v2/swagger.json |
| OpenTrials (CC0) | 2 | https://api.opentrials.net/v1/swagger.yaml |
| Proteomics Identifications Database | 1.2 | http://www.ebi.ac.uk/pride/ws/archive/#!/peptide |
| South African Natural Compounds Database | 1.2 | https://sancdb.rubi.ru.ac.za/docs/ |
| UniProt (CC-BY-ND 3.0 Unported) | 2 | http://www.ebi.ac.uk/proteins/api/doc/swagger/ |
| WikiPathways (CC0) | 2 | http://webservice.wikipathways.org/ |
| XMetDB (CC) | 1.2 | http://xmetdb.github.io/xmetdb-docs/ |

It should be noted that not all of these OpenAPI specification use the same specification version. For eample, the EBI PRIDE (“Proteomics Identifications Database”) database, the XMetDB, the eNanoMapper database, and the South African Natural Compounds Database all use version 1.2 of the OpenAPI specification. Therefore, the other nine OpenAPI specifications use OpenAPI 2 and are suitable for conversion.

### EDAM Ontology Annotation

The ten remaining specifications were annotated with EDAM ontology terms. That annotation covers the input parameters, where *identifier* (edam:data_0842) and *data* (edam:data_0006) are frequently used, but also more specific terms, such as *Ensembl Gene ID* (edam:data_1033). Because most current OpenAPIs are for databases, rather than webservices, a lot of operations are for *Data retrieval* (edam:operation_2422). More detail about the nature of the operation is provided in a hover over:

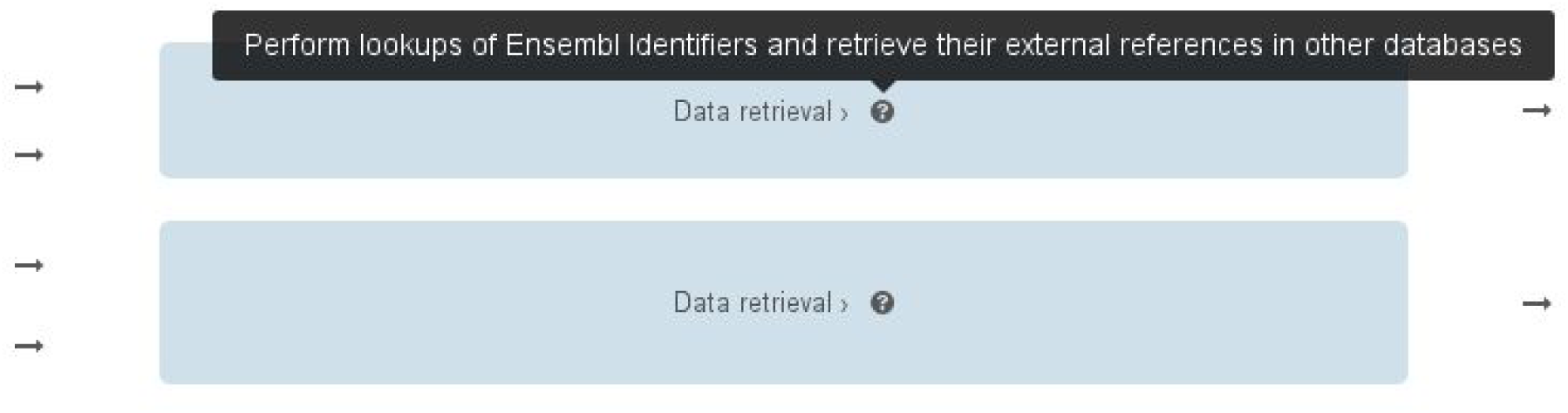

Output is annotated with *Data* (edam:data_0006) or *Identifier* (edam:data_0842), depending on the services. The output formats are also annotated, for example with *HTML* (edam:format_2331), XML (edam:format_2332), and *JSON* (edam:format_3464).

### The convertor

The source code of the convertor is available from GitHub at https://github.com/BiGCAT-UM/swagger2BioTools and with a copy at https://github.com/bio-tools/OpenAPI-Importer. A Maven build file ensure the code can be easily compiled and executed. The code is made available under the MIT license. The code comes with two programs, one that generates an ontology mapping file, *OntologiesMapTemplateBuilder*, using the OpenAPI JSON as input. A second runnable takes that mapping file, along with the JSON file, to output XML suitable for inclusion in bio.tools entries.

### Ensembl OpenAPI

An Ensembl OpenAPI JSON configuration file was generated using the aforementioned approach, resulting in the OpenAPI JSON file available at https://github.com/BiGCAT-UM/EnsemblOpenAPI. This JSON wraps around the REST-like interface provided by the Ensembl database at https://rest.ensembl.org/ [Yates2014]. Various input parameters have been mapped to EDAM ontology terms, in a similar fashion as earlier outlined. For example, the *Ensembl gene ID* is edam:data_1033.

Using the convertor XML was created an uploaded to bio.tools (see link below), resulting in methods visually represented as this:

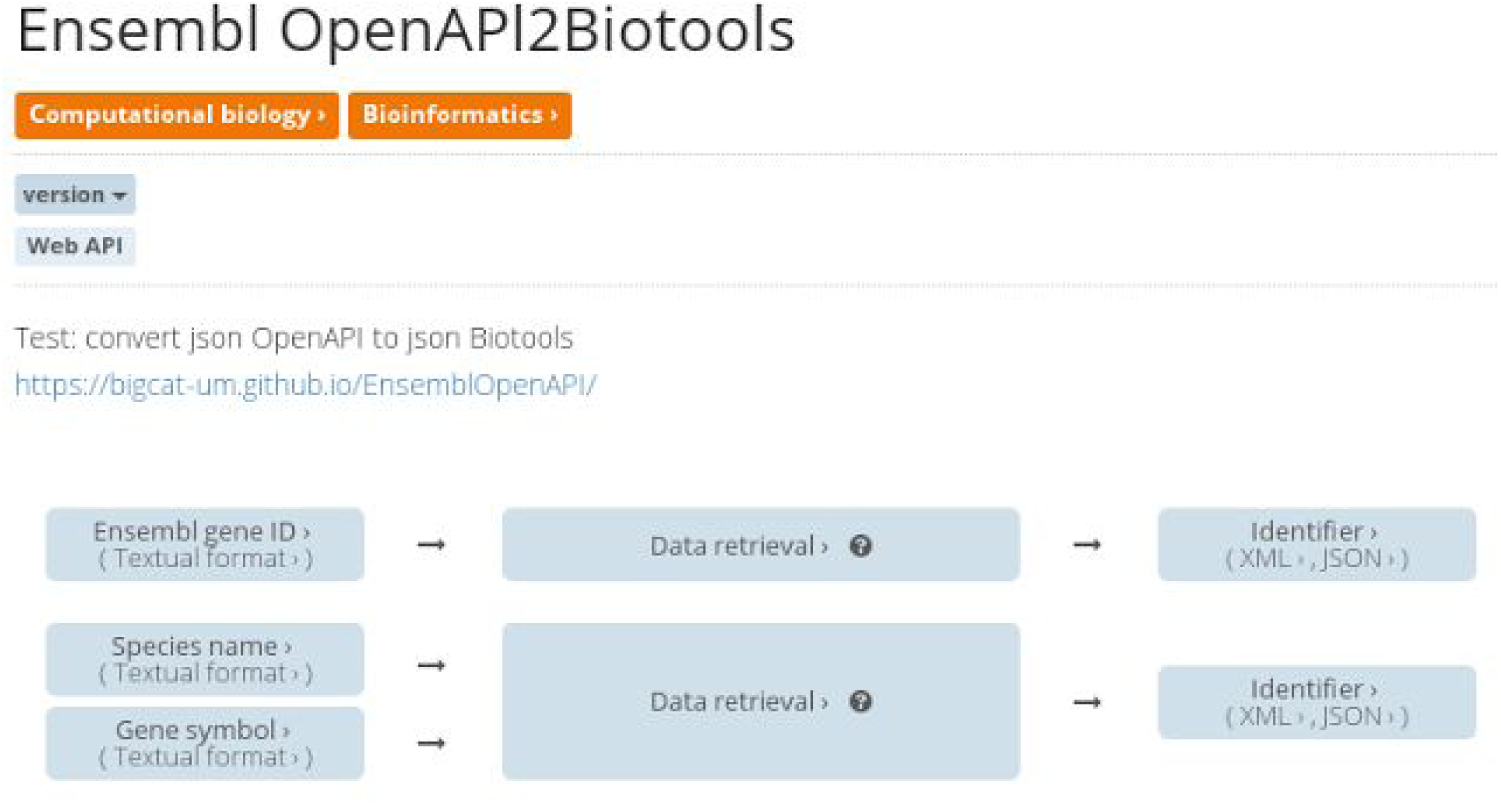

### The bio.tools entries

Taking the OpenAPI JSON created for Ensembl, combined with the JSON configuration for the EDAM ontology annotation of the services, the resulting bio.tools. The below table lists the bio.tools entries that were created.

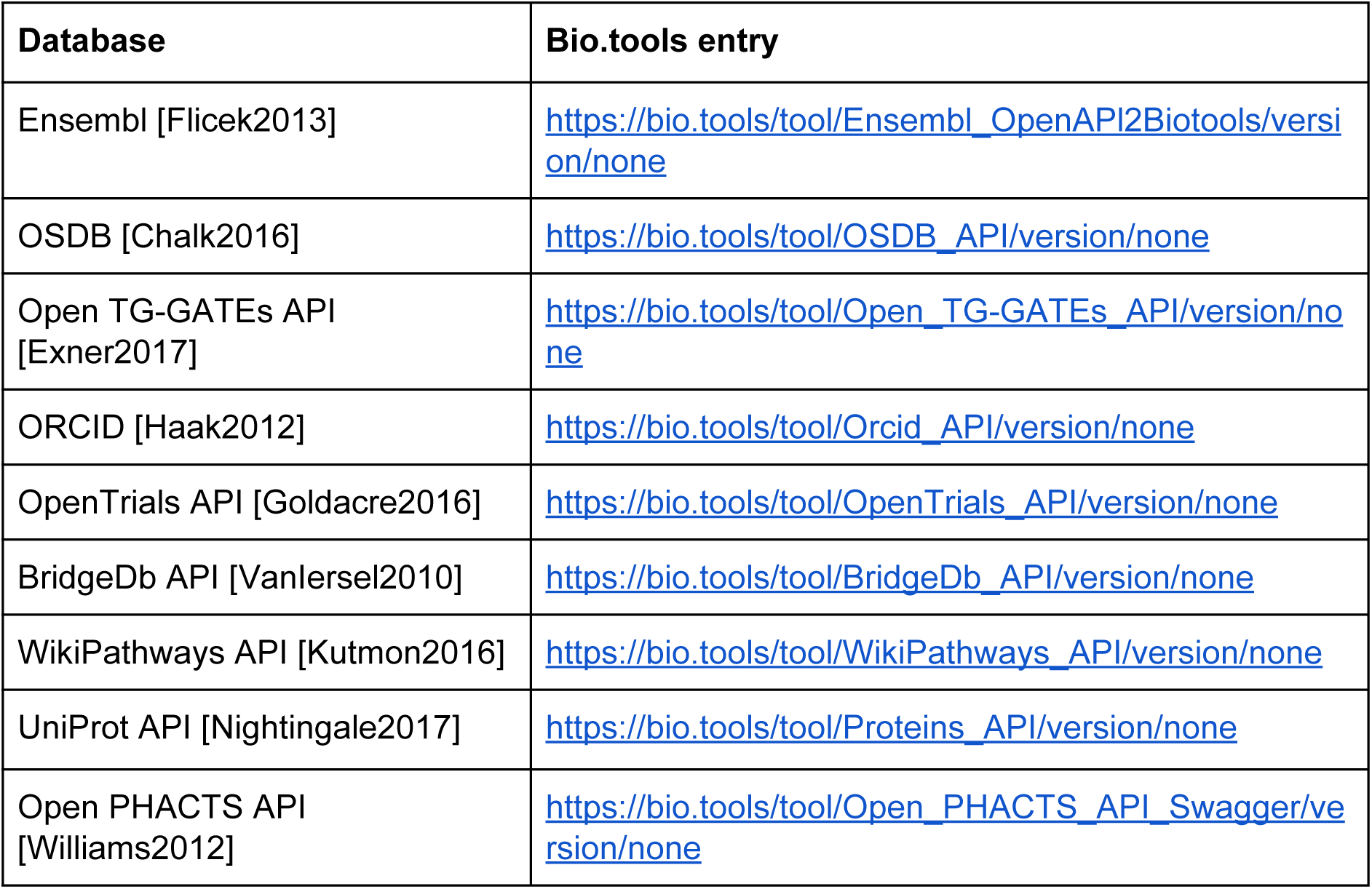

A screenshot of one of the entries (for the OSDB) shows the API entries:

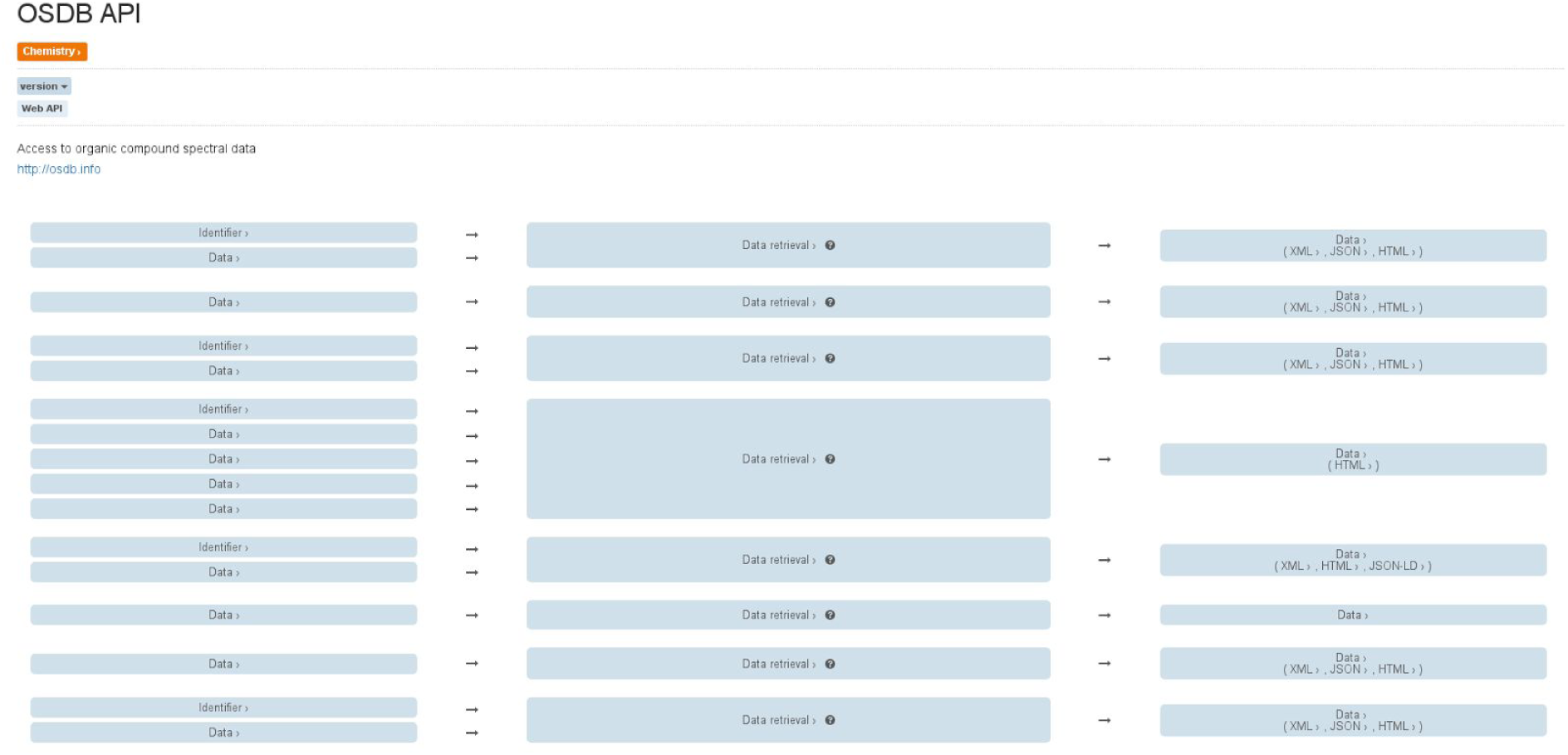

This API only has very general EDAM ontology annotations, though it should be noted that no specific identifier entry is available for some of the supported identifiers, such as the SPLASH [Wolgemuth2016].

## Discussion

The first observation is that the number of publicly available OpenAPI specifications in the life sciences is very disappointing. With hundreds of updated and new databases reported in the database issue of *Nucleic Acids Research*, a meager fourteen webservices is not what I expected. Furthermore, not all of those use the current stable OpenAPI specification, version 2.0, further reducing the number of services for which the convertor works. On the other hand, many more REST and REST-like services exist that can be documented with an OpenAPI specification.

There is plenty of room for improvement. The workflow does not yet take advantage of the API of bio.tools itself. We envision that later versions will inject the service specification JSON directly into the bio.tools entry JSON using this API. An second interesting opportunity is provided by the upcoming OpenAPI 3.0 specification. This new version is more modular, and has the option of specification extensions (https://github.com/OAI/OpenAPI-Specification/blob/OpenAPI.next/versions/3.0.md). This approach can be used to integrate the EDAM ontology annotations directly into the OpenAPI JSON, removing the need for a separate JSON files. One point of attention is that input parameters for web service methods is that they may except different types in the same methods. For example, multiple types of identifiers as input for a single method, where the methods figures out what the type is. In that case, the input of that method needs to be annotated with either a general *Identifier* term, or with multiple terms. However, the technical feasibility of that in bio.tools is yet to be explored. Similarly, we still need to explore if bio.tools allows us to annotate input parameters as optional.

This discussion leads to many new wishes, and, particularly, these specific ideas:

- Support OpenAPI v3 (currently in draft)
- Develop an OpenAPI v3 extension to support ontology annotation
- Align with smartAPI for ontological annotation [Zaveri2017]
- Align with OpenRiskNet [Exner2017]
- Write outreach material about OpenAPI annotations in the life sciences

Support of OpenAPI v3 by existing services is supported by tools that convert OpenAPI v2 configuration files to v3 [Ralphson2017].

## Conclusion

The project has shown how existing and new OpenAPI documentation files can be enriched with ontological annotation and automatically converted into bio.tools entry content. Using this approach, we converted nine OpenAPI JSON configuration files and enriched the matching entries in the bio.tools registry. The number of OpenAPI specifications is unexpectedly low, given the advantages this interactive documentation of webservices offers and the plethora of life science databases published yearly. That said, the upcoming OpenAPI specification is even more powerful, and bio.tools has an opportunity to expert entries as OpenAPI 3.0 to further promote this piece of the FAIR data and interoperability landscape.

## Acknowledgments

We thank ELIXIR-DK for supporting this project via a bio.tools student http://biotools.readthedocs.io/en/latest/studentships.html.

## Appendices Appendix 1: Ensembl

Website: http://www.ensembl.org/

Location of the OpenAPI JSON: https://github.com/BiGCAT-UM/swagger2BioTools/blob/master/resource/input/Ensembl.json

Location of the EDAM ontology annotation JSON: https://github.com/BiGCAT-UM/swagger2BioTools/blob/master/resource/ontology/Ensemblonto.json

Bio.tools entry: https://bio.tools/tool/Ensembl_OpenAPl2Biotools/version/none

## Appendix 2: OSDB

Website: http://osdb.info/

Location of the OpenAPI JSON: http://osdb.info/files/swagger.ison

Location of the EDAM ontology annotation JSON: https://github.com/BiGCAT-UM/swagger2BioTools/blob/master/resource/ontology/OSDB_onto.json

Bio.tools entry: https://bio.tools/tool/OSDB_API/version/none

## Appendix 3: ORCID

Website: https://orcid.org/

Location of the OpenAPI JSON: https://pub.orcid.org/resources/swagger.json

Location of the EDAM ontology annotation JSON: https://github.com/BiGCAT-UM/swagger2BioTools/blob/master/resource/ontology/Orcid_onto.json

Bio.tools entry: https://bio.tools/tool/Orcid_API/version/none

## Appendix 4: Open TG-GATEs

Website: http://toxico.nibiohn.go.jp/english/

Location of the OpenAPI JSON: https://open-tggates-api.cloud.douglasconnect.com/v2/swagger.json

Location of the EDAM ontology annotation JSON: https://github.com/BiGCAT-UM/swagger2BioTools/blob/master/resource/ontology/Tggates_onto.ison

Bio.tools entry: https://bio.tools/tool/Open_TG-GATEs_API/version/none

## Appendix 5: OpenTrials API

Website: https://opentrials.net/

Location of the OpenAPI JSON: https://api.opentrials.net/v1/swagger.yaml

Location of the EDAM ontology annotation JSON: https://github.com/BiGCAT-UM/swagger2BioTools/blob/master/resource/ontology/OpenTrials_onto.json

Bio.tools entry: https://bio.tools/tool/OpenTrials_API/version/none

## Appendix 6: BridgeDb API

Website: http://www.bridgedb.org/

Location of the OpenAPI JSON: http://www.bridgedb.org/swagger/swagger.json

Location of the EDAM ontology annotation JSON: https://github.com/BiGCAT-UM/swagger2BioTools/blob/master/resource/ontology/BridgeDb_onto.json

Bio.tools entry: https://bio.tools/tool/BridgeDb_API/version/none

## Appendix 7: WikiPathway API

Website: http://www.wikipathways.org/

Location of the OpenAPI JSON: http://webservice.wikipathways.org/index.php?swagger

Location of the EDAM ontology annotation JSON: https://github.com/BiGCAT-UM/swagger2BioTools/blob/master/resource/ontology/WikiPathways_onto.json

Bio.tools entry: https://bio.tools/tool/WikiPathways_API/version/none

## Appendix 8: Proteins API

Website: http://www.ebi.ac.uk/proteins/api/doc/

Location of the OpenAPI JSON: http://www.ebi.ac.uk/proteins/api/swagger.json

Location of the EDAM ontology annotation JSON: https://github.com/BiGCAT-UM/swagger2BioTools/blob/master/resource/ontology/Proteins_onto.json

Bio.tools entry: https://bio.tools/tool/Proteins_API/version/none

## Appendix 9: Open PHACTS API

Website: https://dev.openphacts.org/docs/2.1

Location of the OpenAPI JSON: https://dev.openphacts.org/swagger/spec/ops_2_1.json

Location of the EDAM ontology annotation JSON: https://github.com/BiGCAT-UM/swagger2BioTools/blob/master/resource/ontology/OpenPHACTS_onto.json

Bio.tools entry: https://bio.tools/tool/Open_PHACTS_API_Swagger/version/none

